# Automated detection and classification of polioviruses from nanopore sequencing reads using piranha

**DOI:** 10.1101/2023.09.05.556319

**Authors:** Áine O’Toole, Rachel Colquhoun, Corey Ansley, Catherine Troman, Daniel Maloney, Zoe Vance, Joyce Akello, Erika Bujaki, Manasi Majumdar, Adnan Khurshid, Yasir Arshad, Muhammad Masroor Alam, Javier Martin, Alexander G Shaw, Nicholas C Grassly, Andrew Rambaut

## Abstract

Widespread surveillance, rapid detection and appropriate intervention will be critical for successful eradication of poliovirus. With deployable next-generation sequencing (NGS) approaches, such as Oxford Nanopore Technologies’ MinION, the time from sample to result can be significantly reduced compared to cell culture and Sanger sequencing. We developed piranha (poliovirus investigation resource automating nanopore haplotype analysis) to aid routine poliovirus testing of both stool and environmental samples and alleviate the bioinformatic bottleneck that often exists for labs adopting novel NGS approaches. Piranha can be used for efficient intratypic differentiation of poliovirus serotypes, for classification of Sabin-like polioviruses and for detection of wild-type and vaccine derived polioviruses. It produces interactive, distributable html reports, as well as summary csv files and consensus poliovirus FASTA sequences. The reports describe each nanopore sequencing run with interpretable plots enabling researchers to easily detect the presence of poliovirus in samples and quickly disseminate their results.

## Introduction

Following the success of smallpox eradication, great strides have been made towards poliovirus eradication. Vaccine rollout has contributed to a 99% decrease in cases of poliovirus since 1988 (WHO.int accessed 2022-11-22; Figure 1A), however this eradication effort has been hindered by re-emergence of vaccine-derived poliovirus (VDPV). Recent trends indicate an uptick in cases of poliovirus since 2016 (Figure 1B) and since 2020 VDPV has been detected in Africa, Asia, North America and Europe (Figure 1C). There are excellent poliovirus surveillance networks across the world, with the Global Polio Laboratory Network (GPLN) represented in 92 countries. For detection of nascent outbreaks it is important to determine the type of poliovirus circulating in order to implement the appropriate intervention.

**Figure 1.**
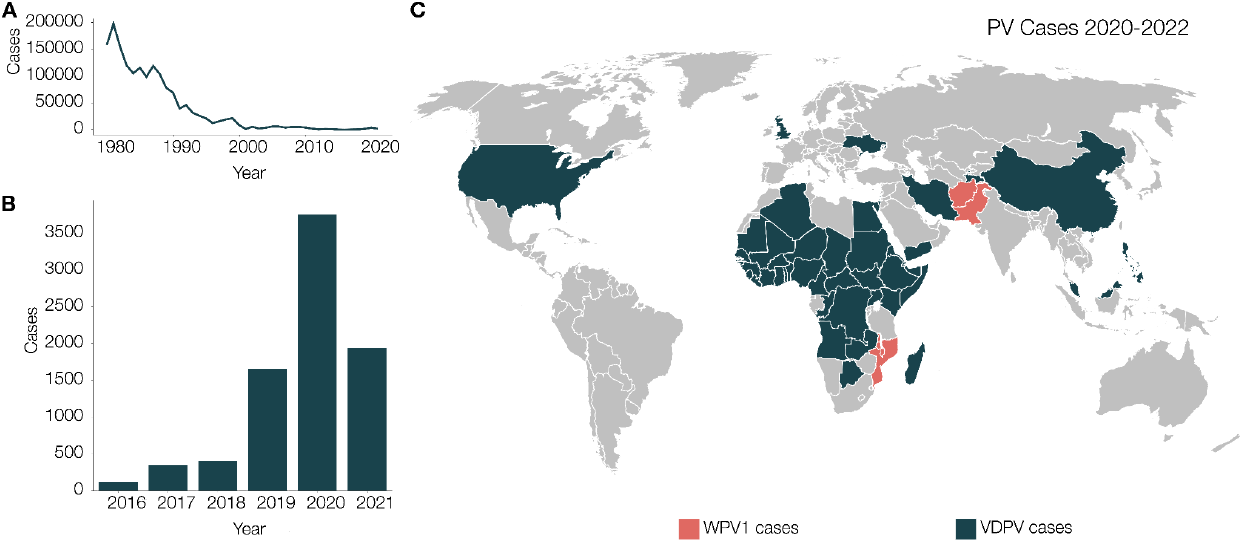
A) Decline in poliovirus cases since 1980. The eradication initiative has helped reduce the incidence of poliovirus by 98% since the 1980s. B) In recent years, there has been an uptick in cases of poliovirus and re-emergence of the virus continues to be a concern. C) Countries in which poliovirus has been detected since 2020. Wild-poliovirus 1 (WPV1) cases indicated in red and circulating vaccine derived poliovirus cases (cVDPV) indicated in blue. cVDPV cases include poliovirus type-1, type-2 and type-3 (VDPV1, VDPV2, VDPV3). Data accessed from WHO.int and polioeradication.org on 2022-11-23. Incidence data from immunizationdata.who.int/pages/incidence/polio.html.

Standard lab algorithms for this characterisation involves culturing stool or environmental samples in poliovirus-susceptible cell lines, intratypic differentiation (ITD) by qPCR and then Sanger sequencing the programmatically important isolates [1,2]. This approach is regarded as the gold standard, however on average it can take 2-3 weeks for cell culturing results. Factoring in transit to a specialised sequencing facility, the median detection time in countries across Asia and Africa ranges from 3-6 weeks [3]. This lag between sample collection and reporting involved with traditional detection approaches has hindered eradication efforts. Shaw et al proposed an alternative detection method that bypasses the need for cell culture and enables direct detection of poliovirus from samples using Nanopore sequencing [4,5]. Nanopore is a long-read technology, which enables us to sequence the VP1 region of poliovirus genomes in a single amplicon [4]. Using this direct detection and nanopore sequencing (DDNS) method to perform sequencing of poliovirus locally within the country of sample testing could result in significant decreases in the time required to detect programmatically important viruses, leading to quicker vaccine responses and truncation of transmission [2].

We are in an era of real-time genomic surveillance. The COVID-19 pandemic illustrated that real-time genomic surveillance is possible and set a precedent for open data sharing practices too. Next-generation sequencing capacity across the globe has grown enormously in the last few years. Whilst bioinformatic capacity has also improved, it is still often a major hurdle to implementing next generation sequencing in-house. The difficulty lies in maintaining a robust and consistent analysis setup, particularly across the entire GPLN where fine scale analysis is critical to determining whether an outbreak is caused by a VDPV or wild-type poliovirus and what should be the appropriate response.

With efficient detection and reporting of poliovirus, governments and health agencies can make informed decisions regarding outbreak control within a timeframe that can help change the course of a poliovirus outbreak, truncating Sabin-related transmission chains before they have an opportunity to re-emerge as pathogenic poliovirus. We present piranha (poliovirus investigation resource automating nanopore haplotype analysis), a novel software tool that provides the means for maintaining consistent, replicable, validated analysis of poliovirus data for both stool sample detection and environmental surveillance.

## Methods

Piranha is a tool developed as part of the Poliovirus Sequencing Consortium (polio-nanopore.org) to help standardise and streamline analysis of nanopore-based poliovirus sequencing, particularly DDNS. It is Python-based with an embedded analysis pipeline built in a Snakemake framework [6]. Piranha can be run as a command line tool or via the piranhaGUI graphical user interface available at github.com/polio-nanopore/piranhaGUI, which handles installation and operating system compatibility problems. The output of piranha includes an interactive HTML report summarising the entire Nanopore sequencing run, a detailed report for each sample within a sequencing run, the output VP1 sequences detected within each sample and variant call support values. Piranha by default is configured to analyse nanopore VP1 sequences generated from the sequencing protocol described in [4]. However, it can be used to analyse whole genome sequences too. It can produce reports in either English (default) or French to aid uptake by laboratories across the GPLN. Documentation describing installation, full usage, configuration options and some example data for piranha is available at https://github.com/polio-nanopore/piranha.

### Input files

Piranha processes nanopore sequencing read files in FASTQ format. Current gold standard for nanopore sequencing analysis is to first basecall and demultiplex raw sequence data using Guppy within MinKNOW [7]. As such, piranha is configured to detect and access FASTQ read data within the specified directory in the format that Guppy produces (subdirectories that correspond to each barcode in the sequencing run with fastq or fastq.gz sequencing read files; Figure 2A). A barcode csv file (Figure 2B) must be supplied, which informs piranha which barcodes to analyse. Minimally, this file must contain information matching a barcode ID (in the same format as the read directory output by Guppy) to a sample ID. Additional metadata can be supplied in this file and it will be incorporated into the final output of piranha to aid in the interpretation of results and data management. For example, in Figure 2B EPID (a unique case identifier) and date have been included as optional metadata columns. There are many configuration options available in piranha, with default settings tailored to the DDNS protocol. All options can be configured with command line flags, or can be supplied in a configuration file in YAML format (Figure 2C).

**Figure 2.**
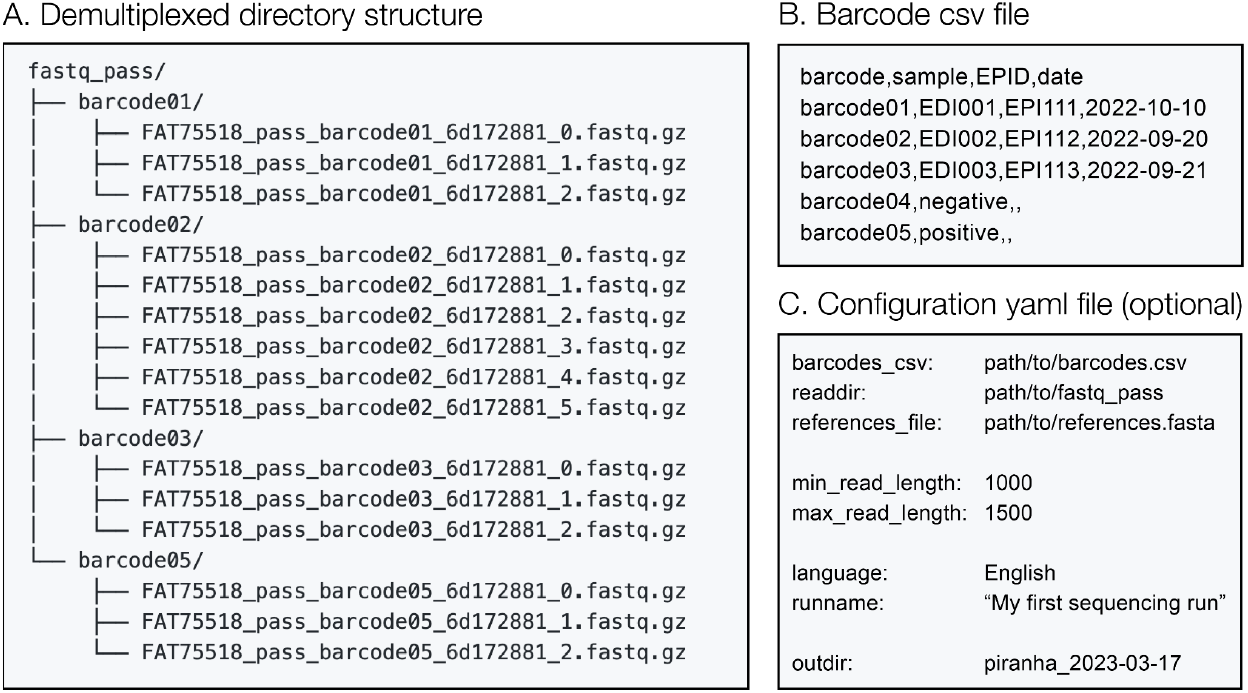
Piranha input files and format. A) Piranha detects fastq (or fastq.gz) files within the demultiplexed read directory specified, in this example ‘fastq_pass’. Within that directory piranha looks for subdirectories with each barcode name and processes all read files within them. Specify the input read directory with `-i/–-readdir`. B) The user must supply a csv file containing minimally a column describing which barcode, and a column mapping a barcode onto a sample ID (`-b/–-barcodes-csv`). Additional metadata can also be supplied, such as in this example EPID or date. C) Users can optionally supply a configuration file (`-c/–-config`)with all command line parameters specified.

### Background database

Within the analysis framework of piranha, noisy nanopore sequencing reads are matched against a reference database using minimap2 (with map-ont mode and no secondary chains reported) [8]. The default reference database in piranha contains a representative set of 959 publicly available sequences for poliovirus type 1, 2, and 3 and non-polio enterovirus VP1 sequences collected from NCBI and VIPR [9](Figure 3; https://github.com/polio-nanopore/piranha/blob/main/piranha/data/references.vp1.fasta). The reference sequences come installed as part of the piranha software, but can also be accessed at github.com/polio-nanopore/piranha. All three Sabin vaccine reference sequences are included in the default database.

**Figure 3.**
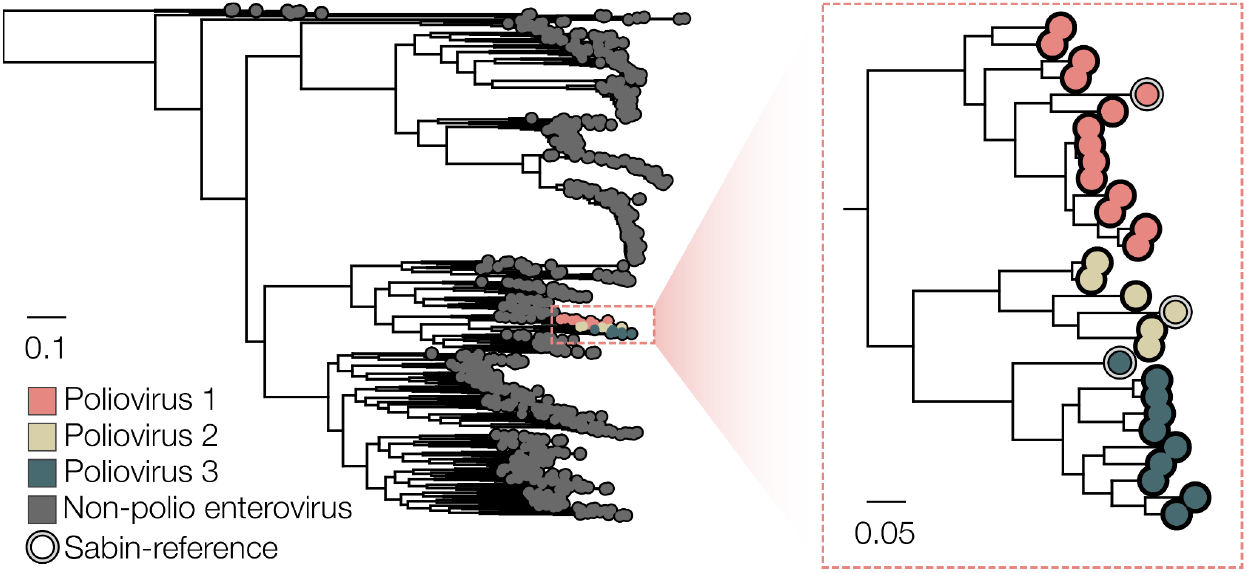
VP1 reference database composition represented as a phylogeny (Distance bar indicates nucleotide substitutions per site). To construct the phylogeny, we aligned the reference sequences using MAFFT [10], visually inspected the resulting alignment and used iqtree2 to estimate a maximum likelihood phylogeny (HKY model) representing the diversity within the background database [11]. The background database consists of a range of human-infectious non-polio enterovirus sequences, representatives of wild-type poliovirus serotypes 1, 2 and 3, and the vaccine reference strains Sabin 1, Sabin 2 and Sabin 3 (n=959).

### Analysis pipeline

Piranha first reads in the barcode.csv file, checks for unique barcode and sample names, and compiles a set of barcodes to analyse. It checks the read path and ensures there are demultiplexed FASTQ files found that correspond to the barcodes it has been instructed to analyse. For each noisy nanopore sequencing read (FASTQs) file, piranha filters by read length to remove any chimeric or off-target reads (Figure 4A). The default length filter is tailored to the VP1 protocol and filters out reads less than 1,000 or greater than 1,500 nucleotide bases long. These defaults can be overwritten by customising the read lengths (`--min-read-length,--max-read-length`) or by using the `--analysis-mode` flag (Analysis modes for pan-enterovirus (`panev`) filters read lengths outwith 3000-4500 and whole genome mode (`wg`) filters for lengths 3400-5200). The length-filtered reads are written to an aggregated read file, one for each barcode specified. This aggregated file is only created transiently unless piranha is run in no-temp mode (specified with `--no-temp`).

**Figure 4.**
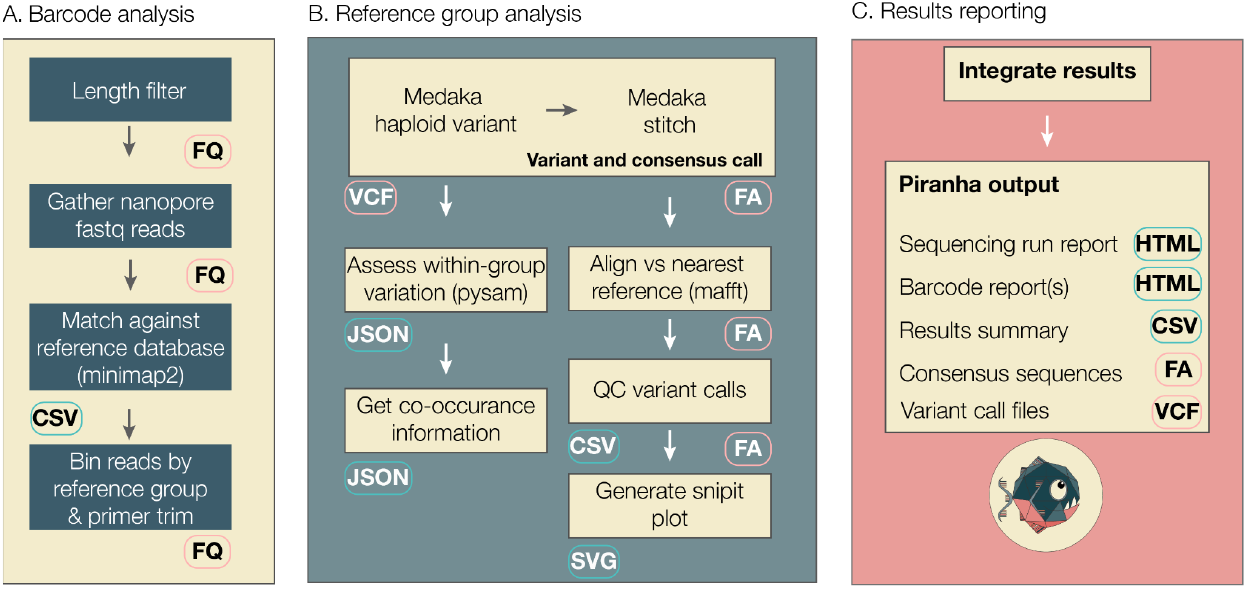
Piranha workflow schema. A) Initial analysis is done for each barcode specified in a barcodes csv file. Reads are mapped against a background reference FASTA file and categorised into broad ‘reference group’ categories: either wild-poliovirus 1 (WPV1), WPV2, WPV3, Sabin1-like, Sabin2-like, Sabin3-like or non-polio enterovirus (NonPolioEV). B) For each reference group detected within a sample, variants are called against the relevant references using medaka and consensus sequences are generated. Within-group variation and co-occurrence information is estimated and QC performed on the variant calls. Snipit plots are also generated based on reference-consensus alignments for each reference group. C) Piranha outputs an interactive summary report and accompanying summary csv, it also produces detailed interactive reports for each barcode and the consensus sequences and variant call files for each reference group detected within a barcode.

Within piranha, each aggregated FASTQ read file is mapped against a FASTA database of reference sequences described in Figure 3. Alternative reference sequences in FASTA format can be supplied to piranha (`-r` flag). Within piranha, these diverse references are categorised into the following bins for further analysis: Sabin1-like, Sabin2-like, Sabin3-like, wild-type poliovirus 1 (WPV1), WPV2 and WPV3 and non-polio enterovirus (NonPolioEV) (Figure 4A). Mapped reads must be within the minimum and maximum read length and have a mapping quality score greater than the minimum mapping quality (default Q=50).

For each barcode, any reference group category that has more than a minimum number of reads and a minimum percentage of sample is taken forward to reference-group analysis (Figure 4B). For each set of reads, piranha first trims the primer sequences (primer length configurable) and then uses ONT’s medaka software to variant and consensus call the noisy nanopore reads against the relevant reference sequence [12]. For Sabin-related reads, they are variant-called against the respective Sabin reference sequence (1, 2 or 3). For wild-type polio and non-polio enterovirus sequences, the reference with the highest number of reads mapping for a given category is used for the variant calling step, resulting in the majority consensus sequence present in a sample for each reference group. Within-group variation is calculated using pysam (https://github.com/pysam-developers/pysam) and we calculate a co-occurrence matrix for variants above the minimum percentage threshold. These thresholds are all configurable parameters either by the command line or through the yaml file. A reference sequence vs consensus sequence is constructed using mafft [10] and biologically implausible variants are replaced with N as they may be a result of sequencing error rather than genuine mutations (e.g. frameshift insertion or deletion mutations in the VP1 coding sequence). Piranha generates snipit plots (https://github.com/aineniamh/snipit) based on the alignments to highlight the differences between reference and sample sequences. This is particularly important for categorising Sabin-related sequences into Sabin, Sabin-like or circulating vaccine-derived poliovirus (cVDPV). For Sabin1-related and Sabin3-related sequences, VP1 sequences with more than 10 mutations relative to the Sabin reference are classified VDPV sequences, whereas for Sabin2-related sequences the threshold is 6 mutations [13].

Piranha produces a number of final results (Figure 4C), by default intermediate workings are created in a temporary directory that gets deleted upon completion but all intermediate files can be retained with the `--no-temp` flag. Piranha outputs the consensus fasta files and variant call files for each reference group present in each barcode, an overall summary csv for the entire run, an interactive html report summarising and visualising the entire sequencing run (Figure 5) and a detailed html report for each barcode (Figure 6). Example reports are hosted at http://polionanopore.org/piranha/report.html.

**Figure 5.**
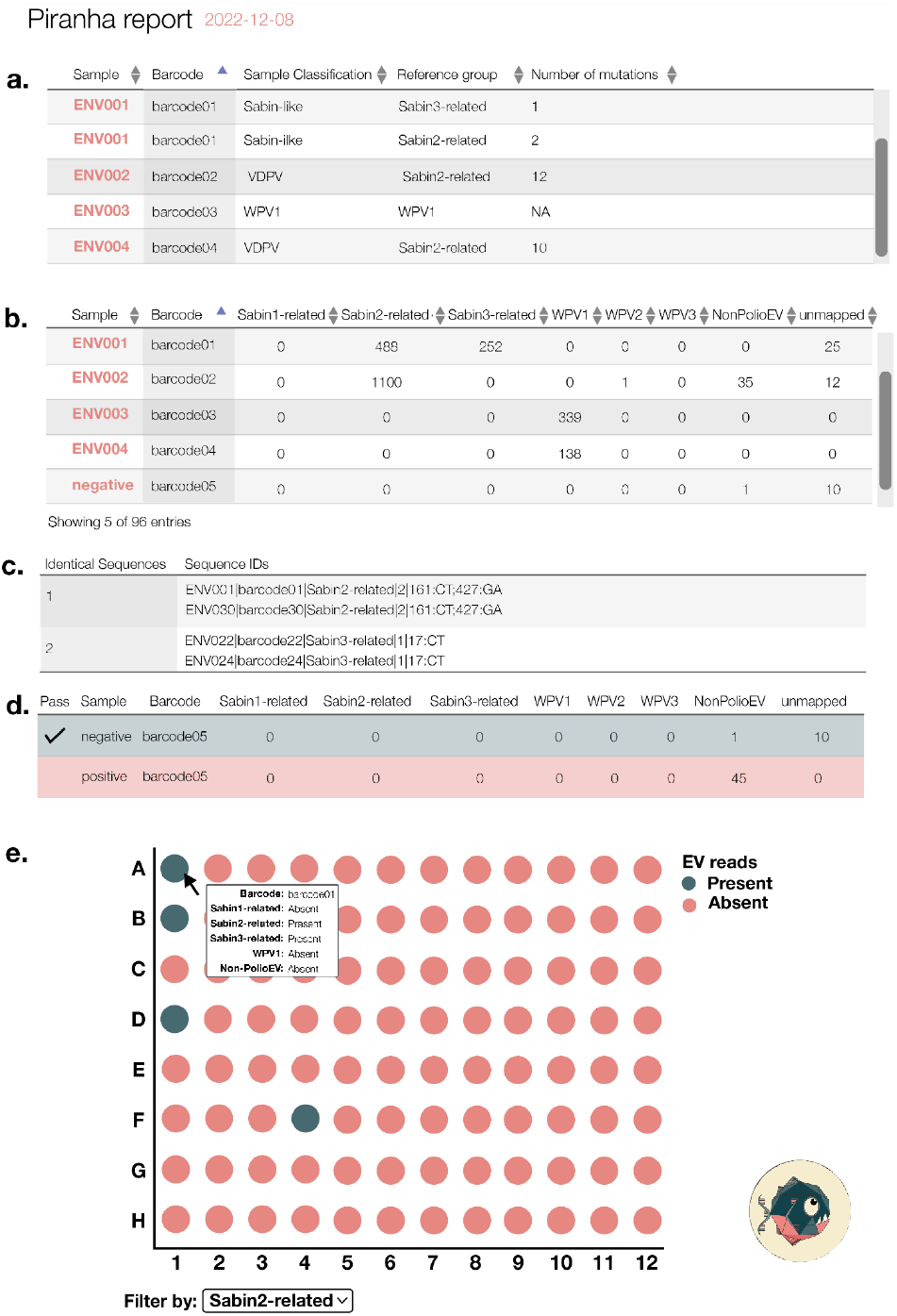
Schema of a piranha output summary report. The report is an interactive, distributable html file and can be personalised, with the ability to add a custom report title, username and institute to the header of the report. A) The first table in the report gives an overview of which enteroviruses were detected in which samples in the sequencing run. For Sabin-related polioviruses, the classification and number of mutations from Sabin are reported. B) The second table includes a summary of the read counts mapped against each reference group for each barcode. C) Any identical sequences present in multiple samples are flagged for investigation as possible contaminants across samples. D) Samples specified as negative or positive controls in the barcodes.csv file are flagged as green with a tick if the control passes, or red if the control fails. For the negative control, samples must have no more than the minimum read threshold mapped in any reference group. For the positive control to pass, samples must have more than the minimum read threshold mapped to the NonPolioEV category since the positive control consists of Coxsackievirus A-20 which is amplified by the DDNS nested PCR. E) The Shaw et al protocol utilises a 96-well plate set up [4]. The piranha report visualises this plate, with barcodes arranged in a configurable horizontal or vertical set up. Presence and absence of different reference groups is indicated to highlight clustering on the plate, again in an effort to flag potential contamination across samples.

**Figure 6.**
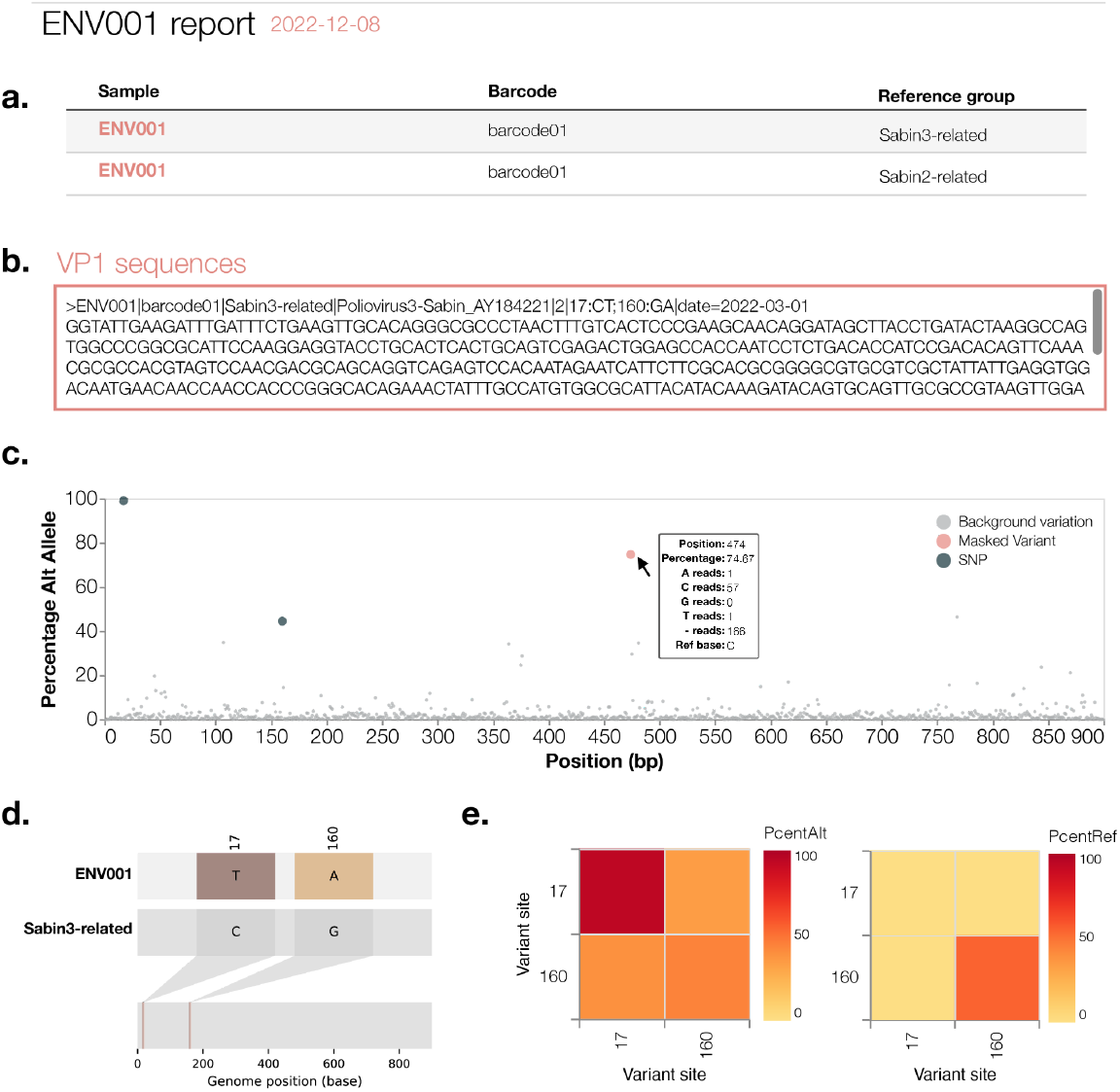
Schema of the detailed piranha report produced for each sample in a sequencing run (here, sample ENV001). A) The first table summarises all reference groups detected in that sample. B) A scrollable box with the consensus FASTA sequences produced for this sample. The sequence record header content is configurable within piranha. C) For each reference group detected, piranha produces a within-group variation plot that shows the percentage of sequencing reads with an allele distinct from the reference allele. The figure highlights the mutations that Oxford Nanopore Technologies’ (ONT’s) medaka called in a given consensus FASTA sequence, and any mutations that may have been masked out by piranha (e.g. a frameshift mutation in VP1). The background % alternative allele variation is displayed in grey. This plot highlights whether there is potential for a mixed sample, as in this example the SNP at position 17 is at 100% frequency, however the SNP called by medaka at position 160 is at ~40% frequency. D) Piranha produces and displays snipit plots, which summarise differences with respect to each reference group detected within a sample. E) Co-occurrence plots summarise the percentage alternative allele and percentage reference for each of the variants called by ONT’s medaka.

## Results

### Piranha can reliably detect Poliovirus1, Poliovirus2 and Poliovirus3 from stool and accurately catalogue the count of SNPs from Sabin for classification of VDPVs

We ran the piranha analysis pipeline on three sets of 200 real nanopore reads generated from sequencing lab-strains of Sabin 1, 2 and 3 respectively, against modified reference sequences derived from Sabin sequences that contain simulated SNPs. The core of the analysis pipeline is medaka haploid variant, which we found to be highly accurate in calling variants in a pure population of reads. We see little to no decline in variant calling ability as simulated SNP count increases for these pure samples until the limit of mapping is reached (Figure 7). At 18.33% divergence we see minimap2 begin to fail to map the reads to the simulated reference. It is then, as alignment accuracy has declined or failed completely that we see variant calls begin to be unreliable. From this finding we can say that within the limits of minimap2’s ability to map reads reliably to a reference medaka should reliably call variants.

**Figure 7.**
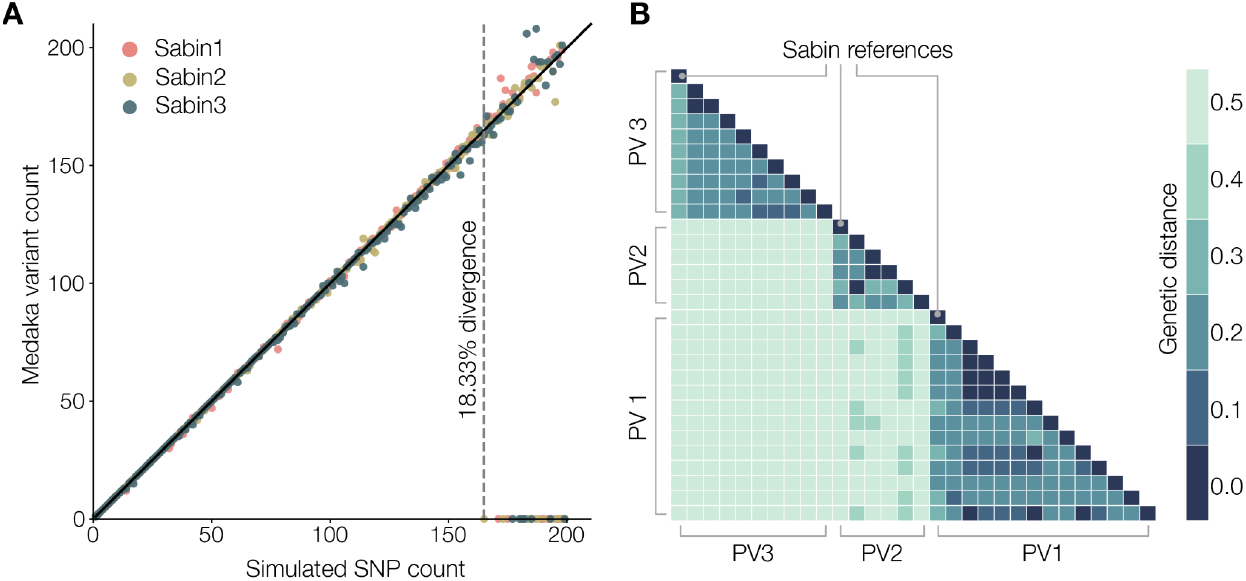
A) We simulated increasing numbers of mutations in a Sabin3 reference away from the actual virus sequence. Variants were called using the medaka haploid variant model with 250 sequencing reads. At 18.33% divergence (165 SNPs from the reference sequence) the reads begin to fail to map using minimap2. B) Pairwise genetic distances (nucleotide substitutions per site) calculated from an alignment of representative Poliovirus (PV) 1, 2 and 3 VP1 sequences. Genetic distance within each Poliovirus group is lower than the observed mapping limit of minimap2.

### Medaka haploid variant found to be most suitable method for variant calling

Within piranha, we wanted to use the most robust variant calling module possible while also considering long term maintenance. Medaka is a tool developed by Oxford Nanopore Technologies that uses neural networks trained on nanopore data to call variants in a sample. Including medaka in piranha’s analysis pipeline was a deliberate choice. Nanopore technology is still in active development and with chemistry and basecalling algorithms changing so frequently, it is important to implement the latest gold-standard analysis suite available for nanopore sequencing. With medaka, updates to the chemistry and upstream analysis algorithms can be co-opted in with the latest trained model. We assessed within medaka which was the most appropriate algorithm for our purposes and found the medaka haploid variant module to be the best at calling SNPs and indels, although it was better at calling SNPs than indels (Figure 8).

**Figure 8.**
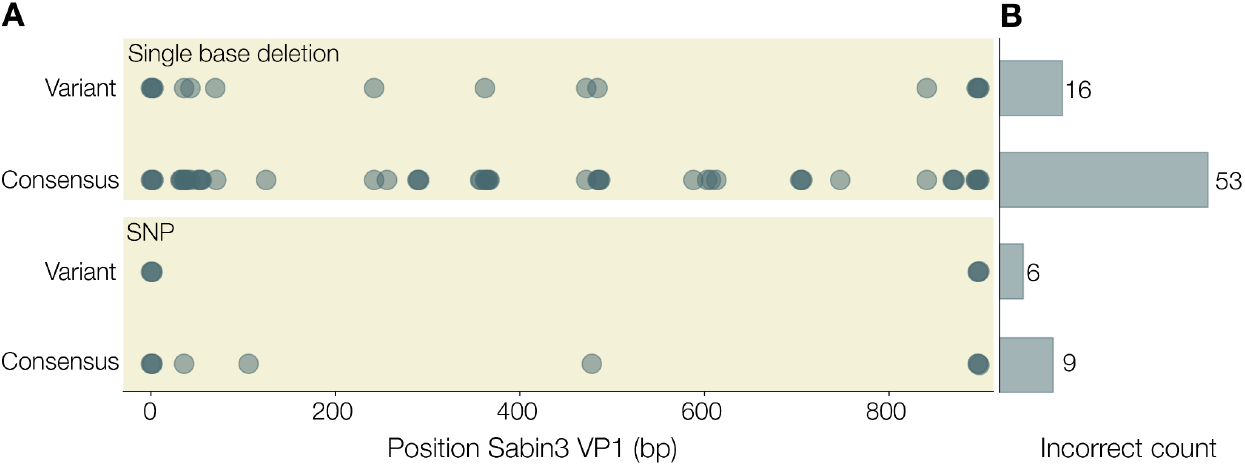
A. We simulated references with either a single base deletion (top panel) or single nucleotide polymorphism (SNP; bottom panel) different from the actual VP1 sequence a set of 250 amplicon reads represent. Points represent incorrect variant calls by either the medaka haploid variant or medaka consensus models of medaka. B. Total count of incorrect variant calls for a given medaka model and type of mutation (deletion or SNP).

### No clear relationship between low-complexity regions and variant quality score

We investigated whether low complexity regions were more likely to give a lower variant quality score and whether variants called at these sites should be reported with the caveat of being within a low-complexity region. We firstly identified low-complexity regions by calculating the Shannon Diversity Index in a 10bp sliding-window across the VP1 region. In all three Sabin reference sequences (1, 2 and 3) we identified a number of low-complexity regions with a score of <1.2 (Figure 9A-C). We confirmed that this index accurately captured low-complexity regions, with regions <1.2 shown in Figure 9D and confirmed these regions captured most homopolymeric regions in the Sabin references (Figure 10). Although there is a correlation between regions of low complexity and variant quality score, the relationship is noisy and likely couldn’t be used to predict the quality of a variant (Figure 11). As such, we provide the variant call files from ONT’s medaka with individual quality scores available for each variant called.

**Figure 9.**
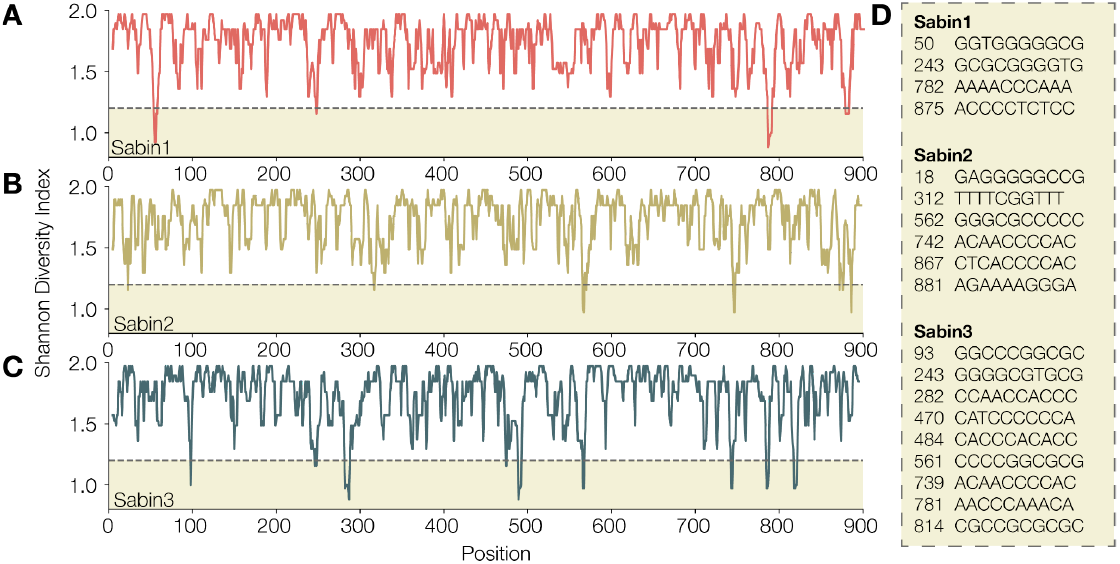
Low complexity regions are highlighted by calculating Shannon diversity index in a 10bp sliding window across the VP1 genes of A) Sabin 1 B) Sabin 2 and C) Sabin 3. Diversity index scores less than 1.2 are highlighted and the VP1 nucleotide position and nucleotides within the 10bp window are indicated in panel D.

**Figure 10.**
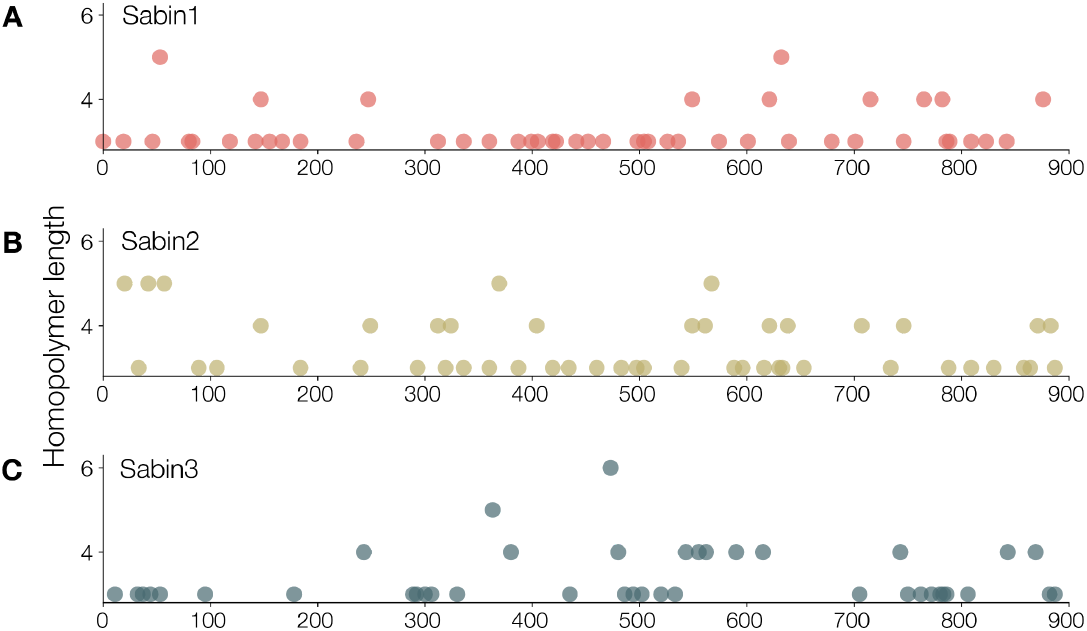
Length of homopolymers across A) Sabin 1 B) Sabin 2 and C) Sabin 3. Nanopore technology struggles with homopolymeric runs. Here we assess the length of homopolymers across the VP1 region for each Sabin reference in turn, to identify regions that may be problematic for variant calling. Homopolymers of length 3 or greater are displayed.

**Figure 11.**
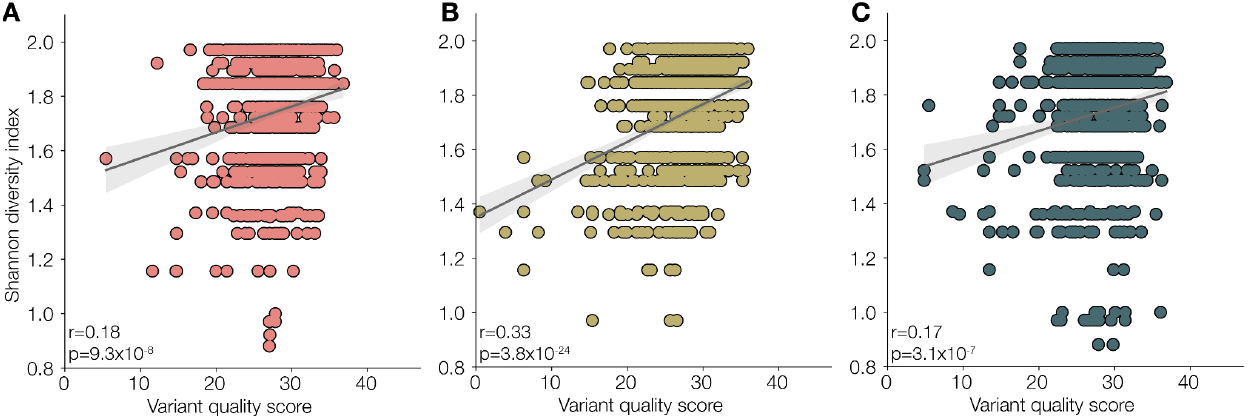
Shannon diversity of nucleotide context index plotted against the quality of the variant call for A) Sabin1, B) Sabin2 or C) Sabin3. Pearson’s r of the correlation and p-value are shown for each plot. The relationship is noisy and variants with particularly low Shannon indexes are not necessarily the lowest quality variant calls.

### Distinguishing between closely related references

We assessed how well we can distinguish between two closely related populations in piranha, which uses minimap2 to map against the reference database. We constructed reference databases of size two with a Sabin1 reference strain and a simulated reference with a fixed number of mutations, X, added to the Sabin1 reference. We simulated these reference databases for X in the range of 1 to 50 (i.e. a series of databases ranging from one containing Sabin1 and a synthetic reference 1 mutation away from Sabin1 and a database containing Sabin1 and a synthetic reference 50 mutations away from Sabin1). For each X, we randomly selected sites to mutate and repeated this 200 times, giving 10,000 test databases each containing 2 references (Sabin 1 and the synthetic reference). We tested how well minimap2 can map the Sabin 1 reads to the correct (Sabin) reference compared with the synthetic reference. We find minimap2 assigns the correct reference with high fidelity (Figure 12). Even at X=1, when the reference sequences are only separated by a single SNP, <1% of reads are assigned the (incorrect) synthetic reference and this stays consistent as the references become more distinct from one another.

**Figure 12.**
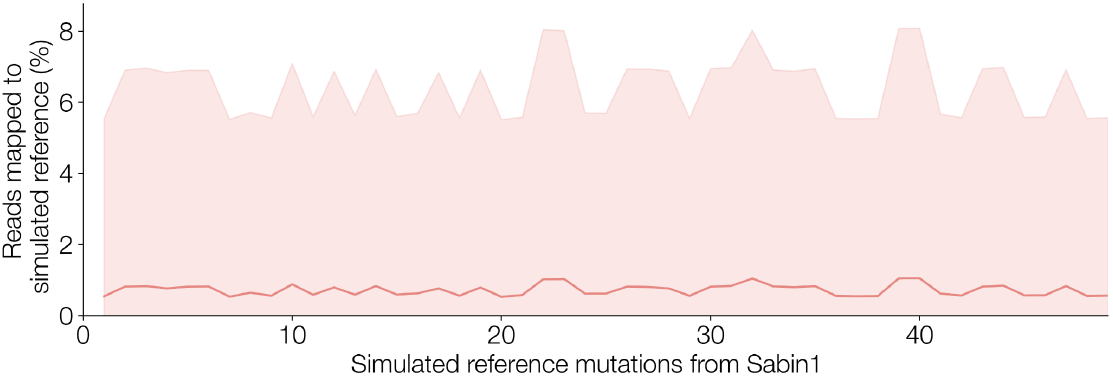
The proportion of Sabin 1 reads mapped to a synthetic (incorrect) reference plotted against the number of mutations separating that reference from the correct Sabin 2 reference. Minimap2 distinguishes between closely related references with very high accuracy. A set of 200 Sabin1 sequencing reads were mapped (-x map-ont) against reference databases containing only a Sabin1 reference and a simulated reference with X number of mutations from Sabin1. For a given number of mutations, we simulated 200 randomly mutated references and so constructed 200 reference files and repeated up to 50 mutations from Sabin 1 (n=10,000). The shaded regions represent standard deviation from the mean values for a given number of mutations, X.

### Distinguishing mixtures that have been created in silico

As a proof of principle, we also confirmed that minimap2 can distinguish between two distinct populations of reads representing different reference groups (in this case Sabin 1 and Sabin 2). Mixtures of real reads were constructed *in silico* at various proportions and minimap2 can resolve these mixtures accurately (Figure 13).

**Figure 13.**
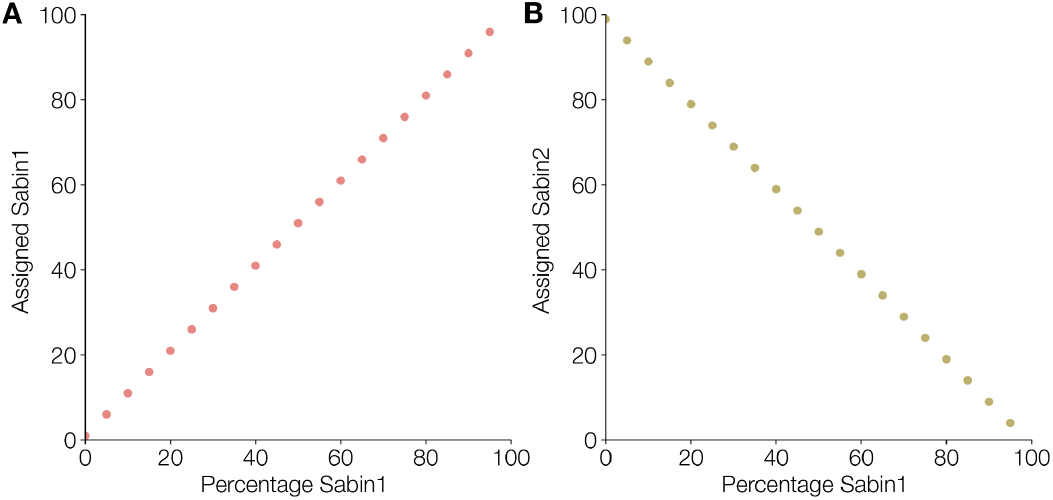
Minimap2 can perfectly resolve mixtures of Sabin1 and Sabin2 sequencing reads constructed *in silico*. As the percentage of Sabin1 in the sample increases, the number assigned increases (A), and similarly the number of reads assigned Sabin2 declines (B).

## Discussion

Despite great strides towards polio eradication, nascent outbreaks arise and continue to persist in many parts of the world. Real-time surveillance and rapid outbreak response will be critical to the elimination of the disease. Deployable technologies for next-generation sequencing, such as the ONT MinION, can aid in rapid outbreak detection and inform interventions weeks earlier than the traditional cell-culture approach [2]. However, these novel methods come with analytical challenges that may be a roadblock for labs considering adopting them. With piranha, researchers have access to best-practice, replicable bioinformatic processing and informative reports making Nanopore sequencing for polio surveillance accessible to labs that don’t necessarily have bioinformatics expertise.

Development of piranha to date has focussed primarily on detection and intratypic differentiation of poliovirus, with emphasis on classifying and identifying circulating VDPV. Default configuration enables piranha to process VP1 sequencing reads and optional flags can be used for processing whole genome sequences. Piranha also constructs a single consensus sequence for each poliovirus serotype or Sabin-related poliovirus detected within a sample, but does illustrate frequency within a sample of each variant, which enables the user to interpret whether sub-populations exist within the data. Future development will extend this analysis to focus on finer-grain mixture resolution within serotypes in an effort to provide further clarity for samples taken for wastewater surveillance. We anticipate the rollout of Oxford Nanopore Technology’s R10.4.1 flow cells and chemistry will aid these efforts significantly by reducing the noisiness of sequencing reads.

Piranha can successfully identify poliovirus sequencing reads in samples for VP1 and whole genome sequencing of poliovirus and distinguish them from related non-polio enterovirus sequences at high specificity. It identifies mixtures of distinct poliovirus serotypes and strains within a sample, potential signs of contamination are flagged, and constructs consensus fasta sequences for each different poliovirus detected. The piranha reports present the output of the bioinformatics analysis in an interpretable, actionable way.

## Data availability

piranha is hosted on GitHub and available under a GNU General Public License v3.0. It is also hosted on Bioconda and has an associated Docker image hosted on Docker Hub.

## Notes

### Competing Interest Statement

The authors have declared no competing interest.

